# Basal Ganglia Neurons in Healthy and Parkinsonian Primates Generate Recurring Sequences of Spikes

**DOI:** 10.1101/2022.05.25.493476

**Authors:** Adriana Galvan, Thomas Wichmann

## Abstract

The spiking activity of basal ganglia neurons can be characterized by summary statistics such as the average firing rate, or by measures of firing patterns, such as burst discharges, or oscillatory fluctuations of firing rates. Many of these features are altered by the presence of parkinsonism. This study examined another distinct attribute of firing activity, i.e., the occurrence of repeating sequences of inter-spike intervals. We studied this feature in extracellular electrophysiologic recordings that were made in the basal ganglia of Rhesus monkeys, before and after they had been rendered parkinsonian by treatment with the neurotoxin 1-methyl-4-phyl-1,2,3,6-tetrahydropyridine (MPTP). Neurons in both pallidal segments and in the subthalamic nucleus tended to fire in repeating sequences, typically 2 ISIs long (i.e., involving three spikes). In recordings that were 5000 inter-spike intervals long, 20-40% of spikes participated in one of many sequences with each ISI replicating the sequence pattern with a timing error of ≤1%. Compared to similar analyses in shuffled representations of the same data, sequences were more common in the original representation of ISIs in the subthalamic nucleus and the external pallidal segment. Induction of parkinsonism reduced the proportion of sequence spikes in the external pallidum but increased it in the subthalamic nucleus. We found no relation between the sequence generation and the firing rate of neurons, and, at most, a weak correlation between sequence generation and the incidence of bursts. We conclude that basal ganglia neurons fire in recognizable sequences of ISIs, whose incidence is influenced by the induction of parkinsonism.

## Introduction

Early electrophysiologic studies of the spiking activities of spontaneously active basal ganglia neurons in primates have focused on summary statistics such as the average firing rate, its variability, or the distribution of inter-spike intervals (ISIs). These statistical measures assume that the occurrence of individual action potentials is independent of others, i.e., that each action potential resets a stochastic process according to which the next action potential is triggered. More recent studies of basal ganglia activity have instead focused on firing behavior that is not governed by a random process, and leads to recognizable firing patterns, such as oscillatory fluctuations of firing, the presence of clusters of short ISIs (‘bursts’), or fractality (1-6). Changes in both, the summary parameters and the parameters describing firing patterns have been identified in animal models of basal ganglia diseases, such as the dopamine-depletion models of primate Parkinsonism produced by injections of the neurotoxin 1-methyl-4-phenyl-1,2,3,6-tetrahydropyridine (MPTP) (summarized in (2)). In such animals, neurons in the subthalamic nucleus (STN), the external and internal pallidal segments (GPe and GPi, respectively), and the substantia nigra pars reticulata (SNr) show a rise in the incidence of burst discharges (2, 7-10), and the emergence of prominent low-frequency oscillatory firing patterns, synchronized across neighboring cells (11, 12). Some of these findings have also been identified in human patients with Parkinson’s disease who underwent brain recording sessions as part of functional neurosurgical procedures (13-16).

Identifying features of firing patterns such as bursts or oscillations in individual neurons implies that the temporal order of ISIs of these neurons is governed by cellular or network dynamics that favor the occurrence of statistically improbable arrangements of ISIs. In the current work, we examine this issue further, by studying whether neurons in the primate basal ganglia show (statistically improbable) exactly repeating patterns of ISIs, and whether the prevalence of such repeated sequences is influenced by dopamine depletion. Similarly repeating patterns of discharge were previously identified in sensory systems (17-23), often occurring in relation to, or representing, the occurrence of specific sensory stimuli. Our study, generated using data from normal and parkinsonian primates, shows that strictly repeating sequences of ISIs are also a feature of spontaneous basal ganglia normal/healthy discharge, and that parkinsonism alters the prevalence and duration of such sequences.

## Methods

### General study design

We conducted electrophysiological recordings of the spontaneous single cell activities in the GPe, GPi, and STN in Rhesus macaques, before and after systemic treatment of the animals with MPTP, inducing a permanent parkinsonian state. The source data are a subset of previously published data (4, 10), including all records that contain at least 5,000 ISIs. The first 5,000 ISIs of such records were analyzed to study the occurrence of repeating patterns of ISIs. Data from individual cells for which longer records were available were also analyzed, to study some of the characteristics of the sequence detection process (as shown in figure 1, below).

**Figure 1:**
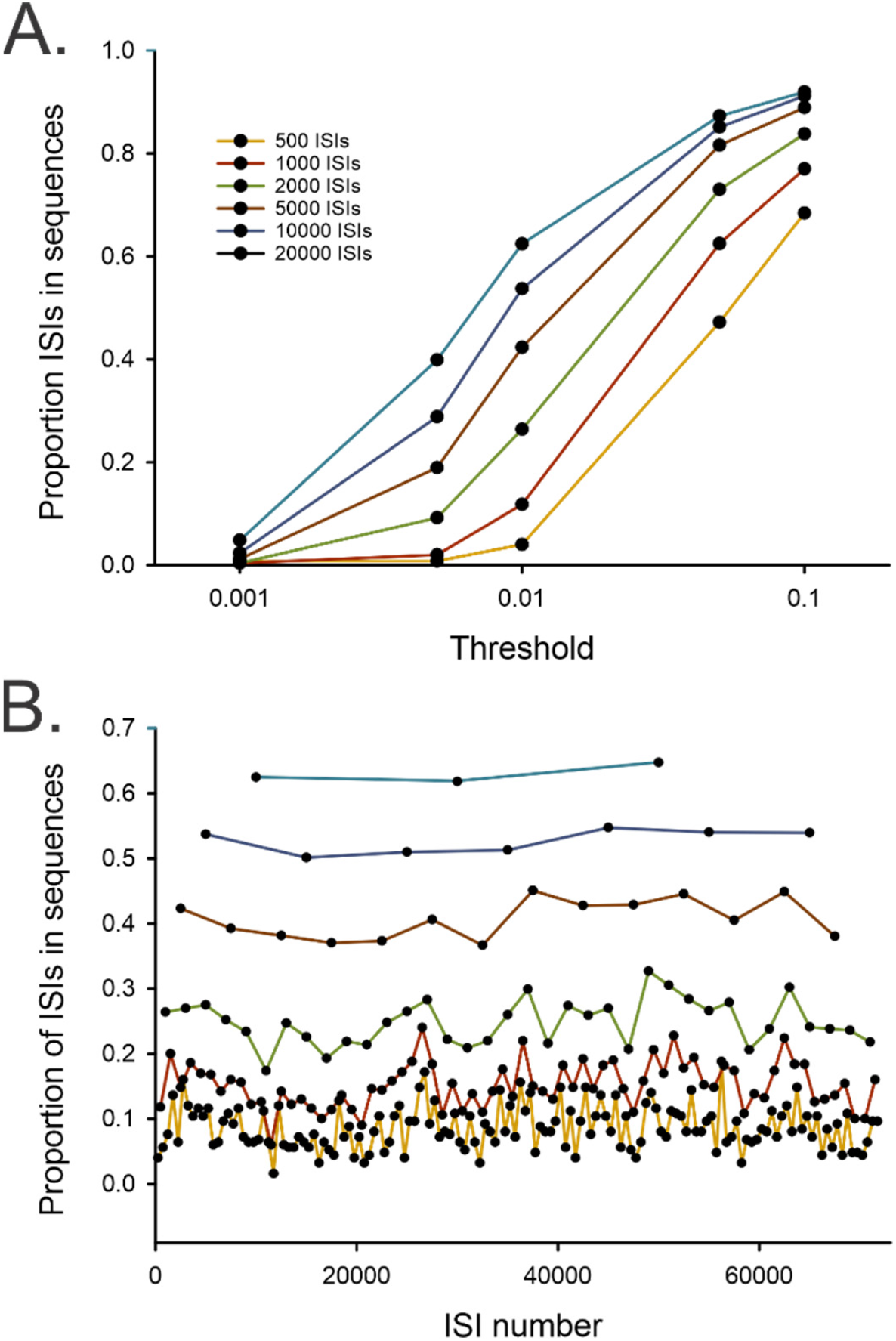
Method of sequence detection. (A) Influence of threshold and length of record on the overall proportion of spikes detected to be contained in sequences as defined in the Methods. The abscissa shows the ‘threshold’ parameter (variability in ISI duration (respective to the ISI length) for an ISI to be accepted in a sequence), and the ordinate the corresponding proportion of spikes identified to be in sequences. The different curves were generated with differing lengths of ISI series (color code indicated in figure), using data from the same neuronal GPe recording. (B) Stability of results of sequence detection. The data were generated using data from a GPe cell that contained more than 70,000 spikes. Data were broken into segments of different lengths, and separately analyzed to yield the proportion of ISIs in sequences (ordinate). Same color code as in A.

### Animals

The data are based on electrophysiologic recordings that were carried out in two Rhesus monkeys (Macaca mulatta, 4-5 kg). The animals were housed under conditions of protected contact housing, with free access to standard primate chow, water, and supplemental fruit and vegetables. Prior to the recording sessions, the animals were adapted to the laboratory environment, and trained to sit in a primate chair and permit handling by the experimenter. All experiments were performed in accordance with the NIH Guide for the Care and Use of Laboratory Animals, the PHS Policy on the Humane Care and Use of Laboratory Animals, and the American Physiological Society’s Guiding Principles in the Care and Use of Animals. All experiments were approved by the Institutional Animal Care and Use Committee of Emory University.

### Surgical procedures

After initial behavioral conditioning, the animals underwent a surgical procedure under aseptic conditions and isoflurane anesthesia (1–3%). Standard stainless steel recording chambers were stereotactically positioned over trephine holes. The animals then received two chambers, one directed at the pallidum (GPe, GPi), placed at a 50° angle from the vertical in the coronal plane, and one directed at the STN, using a 36° angle from the vertical in the sagittal plane. These chambers were affixed to the skull with dental acrylic, along with stainless steel head holders.

### Administration of MPTP and assessment of parkinsonism

After completion of recordings in the normal state, the animals received MPTP. The toxin was injected under general isoflurane anesthesia (1–3%) into the right common carotid artery with the external carotid artery occluded, so that the toxin reached the brain via the internal carotid artery (0.5 mg/kg per injection; one monkey received 2 injections separated by two weeks, while the other received a single injection). Both animals developed similarly obvious signs of moderate parkinsonism (bradykinesia, rigidity, flexed posturing of arm and leg) contralateral to the injections. The animals did not receive any dopaminergic medications. The recordings in the parkinsonian state started 2 months after the MPTP treatment. Throughout the post-MPTP period, the behavioral state of the animals remained stable, as assessed with routine behavioral observations (10, 24, 25). After stable parkinsonism was established, electrophysiological recordings resumed on the animal’s left side (contralateral to the MPTP administration).

### Histology

At the conclusion of the experiments, the monkeys were euthanized by induction of deep anesthesia with an overdose of pentobarbital sodium, followed by transcardial perfusion with saline and 4% paraformaldehyde in 0.1 M phosphate buffer (pH 7.2). The brains were removed and cryoprotected in 30% sucrose solution in 0.1 M phosphate buffer. The fixed brains were sectioned in coronal planes (50 µm). One of every four sections was stained with cresyl violet for localization of microelectrode tracks. The results of the histological examinations are documented in our previous publication (10), showing the placement of the recording electrodes in the basal ganglia, minimal overall damage from the electrode penetrations, as well as the MPTP-treatment induced striatal dopaminergic denervation, based on immunohistochemistry.

### Electrophysiologic recordings

The electrophysiologic recordings were done with the animal seated in a standard primate chair, with its head restrained. Recordings were conducted during episodes when the animal was fully awake, as verified by direct observation. The neuronal activity of basal ganglia neurons was recorded with commercial tungsten microelectrodes (Frederick Haer Co., Bowdoinham, ME; impedance 0.5-1.0 MΩ at 1 kHz). The electrodes were lowered into the brain with a microdrive (MO-95B, Narishige, Tokyo, Japan), using a guide tube. The electrical signals were amplified (DAM-80 amplifier, WPI, Sarasota, FL), filtered (400-10,000 Hz, Krohn-Hite, Brockton, MA), displayed on a digital oscilloscope (DL1540, Yokogawa, Tokyo, Japan), made audible via an audio amplifier, and recorded as digital signals. Neurons in the GPe, GPi, and STN were identified based on their stereotaxic location within the brain, the regional relationship to nearby nuclei, and their characteristic firing patterns, including the findings of neuronal firing at high-frequency with pauses in the GPe, the finding of high-frequency discharging cells without pauses in the internal pallidal segment, and the finding of cells firing at lower rates embedded into an area of high background activity in the STN (see, e.g., 26, 27). The location of neurons was subsequently reconstructed based on stereotaxic information, micromanipulator readings during the recordings, and the results of postmortem histologic analysis. For initial inclusion into the analysis, cells had to be adequately isolated throughout the record, as defined by a signal-to-noise ratio of at least 3. The recorded activity was later replayed and processed with a template-matching spike sorting device (Alpha-Omega, Nazareth, Israel) which extracted the timing of spike occurrence. The data were then stored as series of ISIs.

### Data analysis

The ISI data were imported into the Matlab computing environment (Mathworks, Natick, MA) for further analysis. Using a custom-written Matlab algorithm, we analyzed whether repeating sequences of two or more ISIs could be found in the data stream. A sequence of ISIs was said to repeat itself if the duration of each member of the repeated sequence was within 1% of the corresponding value within the original sequence. In a subsequent step of the algorithm, we examined whether the identified sequences were overlapping or fully contained in one another. Sequences that were fully contained in longer ones were removed. If some repetitions of a short sequence were found to be contained in a larger one, while other repetitions occurred in isolation, the isolated occurrences were retained if they were found more than once. If some instances of the detected sequences overlapped with others, the longer of the overlapping sequence was retained in the record, while the remaining member(s) of the shorter overlapping sequence(s) were discarded.

We considered that any (random) arrangement of ISIs from a given distributions may lead to a finding of repeating sequences. To determine the probability of finding repeating sequences as a random process, given a neuron’s ISI distribution, we applied the steps of identifying sequences (as described above) to 100 randomly arranged representations of the same ISI data for each cell.

The data based on the original ISI stream and those generated from each of the shuffled representations of a cell’s data were used to calculate the total number of sequences, the distribution of sequence length, the distribution of the maximal number of repetitions per sequence length, the proportion of spikes and time spent in sequences, and the median duration of sequences. The results based on the original data were also expressed in proportion to the results from the shuffled data (see figure 3, below).

Given the possibility that the finding of bursts may overlap with the finding of sequences, we were interested in examining how the incidence of sequences corresponded to the occurrence of bursts. We identified the presence of bursts using the ‘surprise’ algorithm (28), using a cutoff-surprise value of 3, allowing us to identify those ISIs that were part of bursts. We then determined the proportion of sequence ISIs that were also identified as burst ISIs, and the proportion of burst ISIs that were also found within sequences. Regression analyses were performed to investigate correlations between the incidence of bursts and that of sequences across the population of neurons.

### Statistics

For each record, we used the paired non-parametric Wilcoxon test to compare the original proportion of spikes in sequences and the proportion of time spent in sequences with the median of the results based on shuffled data. These analyses were done separately for the pre- and post MPTP data. Comparisons between the pre- and post-MPTP data relied on (non-paired) non-parametric Mann-Whitney tests. For all tests, a criterion of p < 0.05 was taken as a measure of significance.

## Results

### Database

The number of cells presented in this report, and the average duration of the records is shown in the table. Sufficient data from a total of 78 neurons from GPe (38 recorded in the normal state, 40 recorded in the MPTP-treated state), 67 neurons from GPi (33 recorded in the normal state, 34 recorded in the parkinsonian state), and 40 neurons from STN (13 recorded in the normal state, 27 recorded in the parkinsonian state) were available. The table also provides information about the firing rate, and the coefficient of variation (CV) of the ISIs. We found that the cells recorded in parkinsonian state in the GPe were significantly slower than those recorded in the normal state. The neurons in the STN fired significantly faster. The firing rate in GPi in the MPTP-treated state was higher than that in normal state, but this difference was not significant. The average coefficient of variation (CV) of ISI durations did not differ between the two states in any of the three regions.

**Table.** Basic firing characteristics of neurons in GPe, STN and GPi.

| Structure | State | Number of ISIs | Duration (s) | Average firing rate (spikes/s) | CV of ISIs |
| --- | --- | --- | --- | --- | --- |
| GPe | Normal<br>(n = 38) | 32,114.2 ± 16,154.9 | 482.1 ± 189.5 | 65.7 ± 19.0 | 1.15 ± 0.46 |
|  | MPTP<br>(n = 40) | 21,989.7 ± 12921.0<br>(p = 0.002) | 459.4 ± 172.8<br>(p = 0.353) | 48.5 ± 19.3<br>(p < 0.001) | 1.30 ± 0.46<br>(p = 0.187) |
| STN | Normal<br>(n = 13) | 11,327.7 ± 3,934.7 | 512.6 ± 111.5 | 22.7 ± 8.2 | 1.45 ± 0.27 |
|  | MPTP<br>(n = 27) | 19,971.0 ± 7,435.6<br>(p < 0.001) | 527.1 ± 125.7<br>(p = 0.613) | 38.2 ± 11.4<br>(p < 0.001) | 1.38 ± 0.36<br>(p = 0.378) |
| GPi | Normal<br>(n = 33) | 36,722.4 ± 14049.9 | 503.2 ± 155.7 | 73.5 ± 18.1 | 0.90 ± 0.22 |
|  | MPTP<br>(n = 34) | 38,779.7 ± 14671.1<br>(p = 0.821) | 495.2 ± 138.2<br>(p = 0.334) | 78.1 ± 17.9<br>(p = 0.444) | 1.03 ± 0.36<br>(p = 0.156) |

*n*-values refer to the number of neurons available for study in each category. *p*-values are based on Mann-Whitney tests, comparing the respective data from the normal and parkinsonian states. See test for abbreviations.

### Results of sequence analysis

The detection of sequences of ISIs in the recorded material was strongly influenced by the threshold chosen for detection of such sequences, as demonstrated in figure 1A, based on data from a single GPe neuron recorded in the normal state for which a long (72,182 spikes, recorded over 842.3 s) record was available. It can be seen that a sequence acceptance threshold of 0.001 (that is, acceptance of ISIs to be in sequences only if their ISI durations differed by less than 1/1000 of the respective ISI lengths) resulted in very low levels of detection, quantified as overall proportion of ISIs in sequences, among all recorded ISIs, while use of a threshold level of 0.1 resulted in a very high proportion of ISIs in sequences. For this study, we settled on an intermediate threshold level, 0.01 (i.e., ISIs were accepted as belonging to the same sequence, if their duration differed by less than 1/100 of the corresponding ISI length in the original sequence).

Figure 1A also demonstrates that the length of the record mattered. As expected, analysis of long data segments resulted in the identification of more sequences (and thus, a greater proportion of spikes in sequences) than analysis of short data segments, likely because repeating ISI sequences that were separated in time can be detected in long records, while they may elude detection in short records. As noted above, we analyzed records with at least 5,000 ISIs. If more than 5,000 ISIs spikes had originally been recorded, we used the initial 5,000 ISIs for the analysis.

The data in figure 1B (same color scheme) demonstrate the stability of the sequence detection across time. This analysis was based on the same record as figure 1A. The record was repeatedly analyzed, breaking it into non-overlapping consecutive segments of equal lengths, ranging from 500 and 20,000 ISIs, calculating the (measured) proportion of spikes in sequences for each. There was considerable variability/uncertainty when measurements involved short sequence lengths (500-2,000 ISIs), while longer data segment lengths (5,000-20,000) yielded more stable results.

Figure 2A demonstrates the results of detection of a specific three-spike sequence of spikes in a GPe cell recorded in the normal state, different from the cell shown figure 1. The sequence was found 9 times in this neuron’s 5,000 ISI data. Such 3-spike sequences (involving 2 ISIs) were, by far, the most common type of sequence identified. Figure 2B shows the number of unique sequences identified across cells in GPe, STN and GPi. Red symbols represent data from neurons recorded in the normal state, while gray symbols represent data from neurons recorded in the parkinsonian state. Few cells showed 4-spike (3-ISI) or 5-spike (4-ISI) sequences. With the induction of parkinsonism, the overall proportion of spikes within sequences changed, as shown in figure 2E. These recordings from STN cells were taken during the normal and parkinsonian states and demonstrate a parkinsonism-related substantial increase in the number of ISIs that were part of sequences. The cell recorded in the parkinsonian state also showed an increase in burst firing and oscillatory activities, but as noted below, we did not find a strong correspondence between these phenomena and the tendency of the cell to generate sequences of firing.

**Figure 2:**
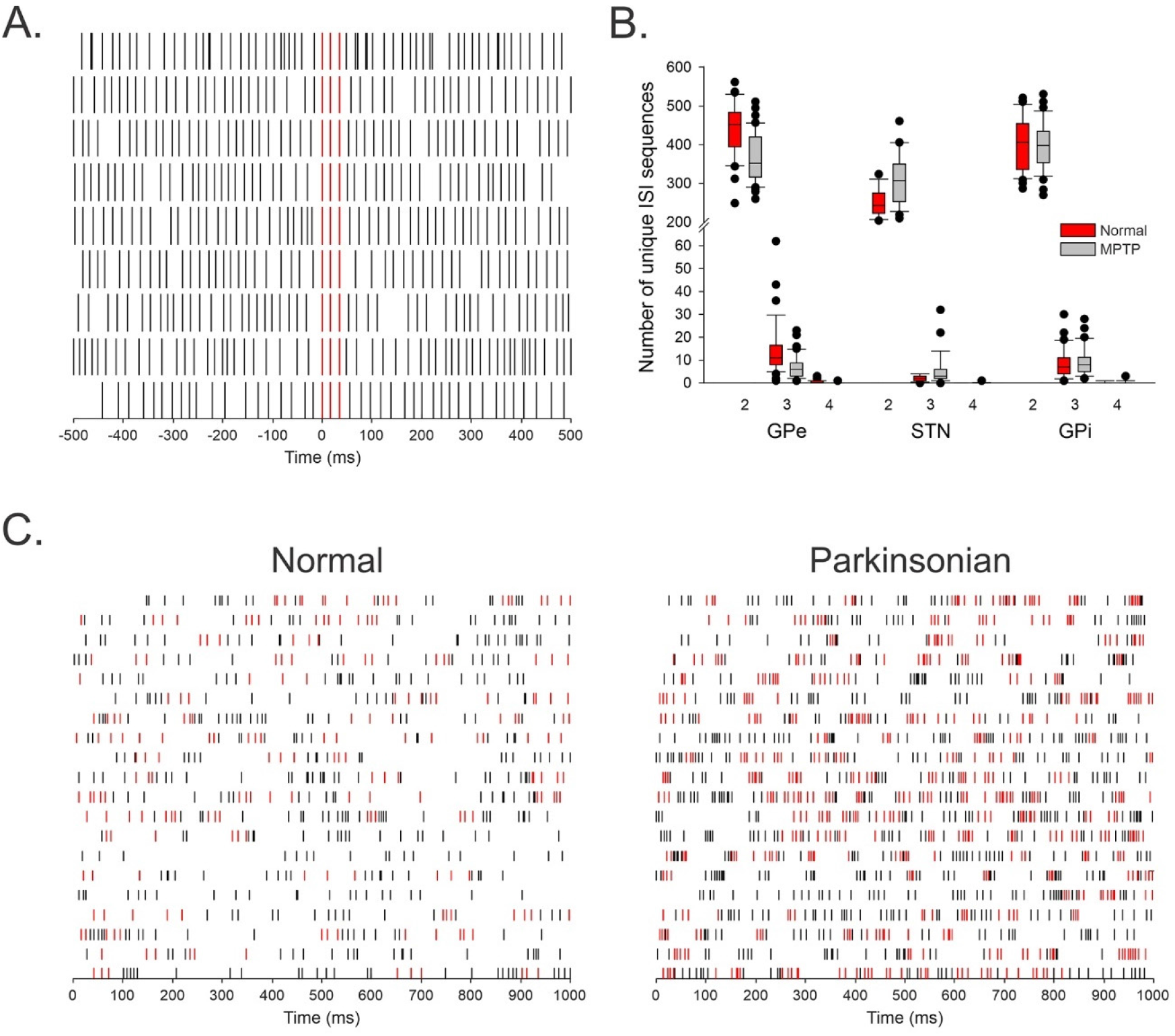
Sequence detection in basal ganglia of normal and parkinsonian monkeys. (A) Example of repeated sequences of 2 ISIs (3 spikes) in a GPe neurons, recorded in a normal animal. Each of the short vertical lines corresponds to a single action potential. The identified sequences were aligned to the first spike in the sequence (sequence indicated in red). (B) Overall distribution of unique sequences of different lengths. The boxplot shows the median number of unique sequences (with 25-75^th^ percentile boxes, 10^th^ and 90^th^ percentile line indicators, and dots indicating values outside of the 10^th^/90^th^ percentile boundaries) comprised of 2, 3 or 4 ISIs in the normal (red) and parkinsonian (gray) state in GPe, STN, or GPi. (C) Examples of 20 second segments, split into 20 consecutive 1s segments, of spiking activity of an STN cell recorded in the normal state (left), and an STN cell recorded in the parkinsonian state (right). The spikes marked in red were parts of sequences recorded in these cells (based on analysis of the first 5000 ISIs of the neuron, see Methods).

### Proportion of ISIs participating in sequences

An analysis of the proportion of cells participating in sequences is presented in figure 3, with statistical details provided in supplementary table 1. The left panel of fig. 3A shows a comparison of the proportions of spikes in sequences in cells recorded in the normal and parkinsonian states. The tendency to fire in sequences was almost twice as high in the pallidum than it was in the STN in the normal state. After induction of parkinsonism, this proportion dropped significantly in the GPe cells, while it increased significantly in neurons recorded in the STN. There was no change of this parameter in the GPi recordings. In the right panel of fig. 3A, we compared the proportions of spikes found in sequences with the median of the proportion of spikes in sequences based on shuffled representations of each cell. In most cases, we found that the proportion of sequences in the recorded arrangement of ISIs was significantly more common than in their shuffled representation suggesting that the cellular activity favored the occurrence of sequences. This was the case for ISIs in GPe and STN in the normal and parkinsonian states, shown by the red and gray box plots, respectively. In GPi, the finding of sequences in the normal state is most likely explained as a stochastic phenomenon because there was no statistical difference to the finding of sequences in the shuffled representations. However, post-MPTP, more sequences were found in GPi cell recordings than in random arrangements of the same ISIs.

**Figure 3:**
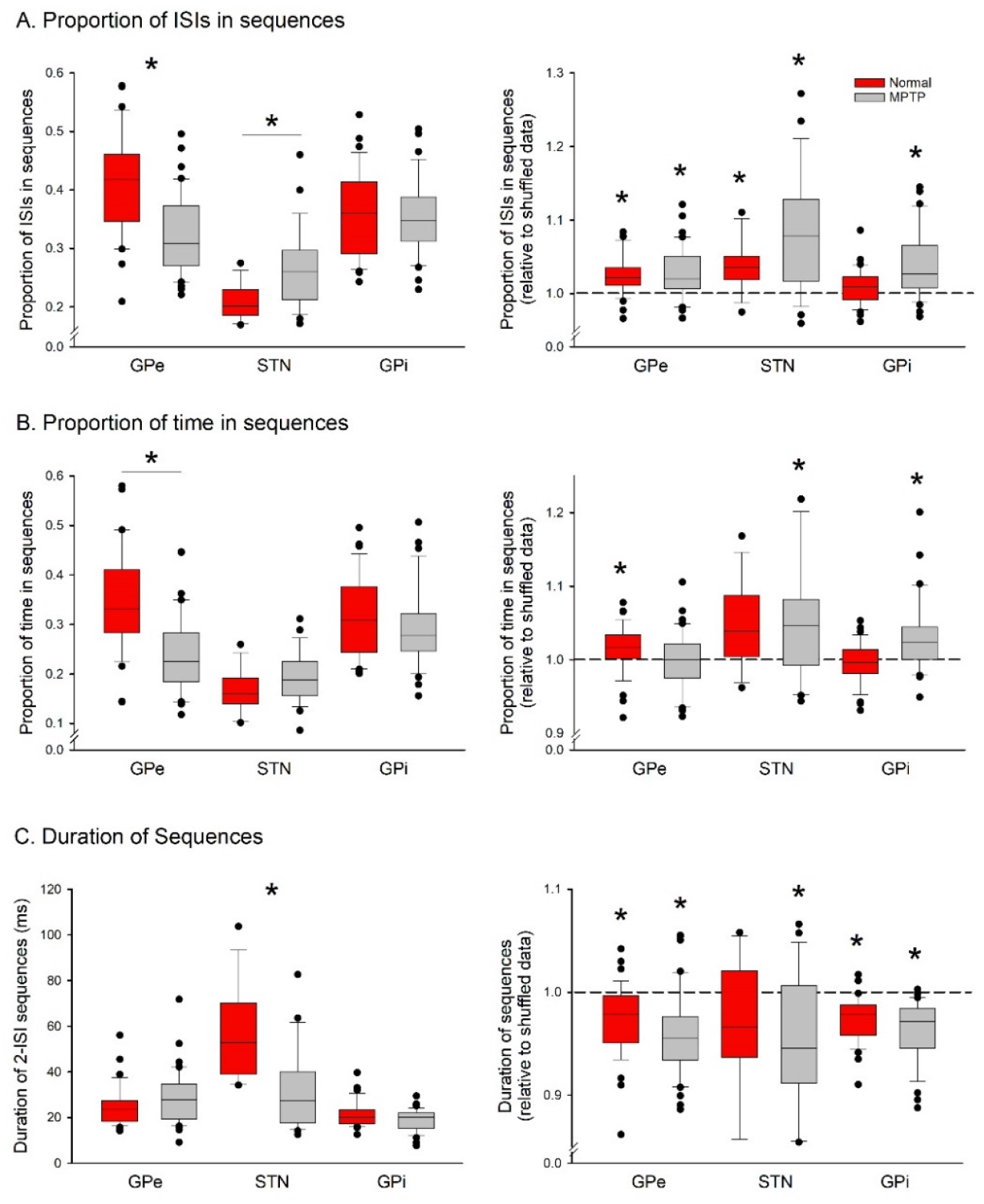
Proportion of ISIs in sequences (A), proportion of time spent in sequences (B), and duration of 2-ISI sequences (C). The box plots in the left panels shows the median of the respective absolute values. The graphs on the right show the distribution of the ratios between the original data and the respective median of 100 shuffled renditions of the same ISI data. Cells in GPe, STN and GPi were recorded in the normal state (red) and in the parkinsonian state (gray). The boxes show the 25^th^ and 75^th^ percentile ranges, with the median marked inside of the box, the error bars show the 10^th^ and 90^th^ percentile, and the black dots indicating values outside of the 10^th^/90^th^ percentile boundaries. Sequences were defined as described in the text. The analysis is based on the first 5,000 ISIs in each data file. *, p < 0.05; Mann-Whitney test was used for plots on the left, comparing the data from the normal and parkinsonian states. The paired Wilcoxon test was used for plots on the right, comparing the median duration of sequences of the actual data with that of the median of sequence analysis of the shuffled renditions for each cell.

An analysis of the time spent in sequences (shown in figure 3B) revealed results similar to those found in figure 3A. In figure 3C, the duration of sequences is compared. As this measure would obviously differ between sequences containing different numbers of spikes, we restricted this analysis to the most common type, i.e., sequences containing 3 spikes (see also fig. 2B). This analysis showed that sequences had similar length in the GPe and GPi samples from the normal and parkinsonian state but were longer in duration in the STN in the normal state than in the parkinsonian state. Comparison with shuffled data showed that the sequences were shorter in the recorded data in GPe and GPi in both, the normal and parkinsonian states, and in the STN data in the MPTP-treated state. This suggests that, in almost all groups, one or both ISIs in the sequence data was shorter than the median ISI length.

### Relationship between the proportion of spikes in sequences, the firing rate, and bursts

We explored the relationship between sequences and the frequency of firing, and the occurrence of bursts with the analysis shown in fig. 4. The proportion of spikes in sequences was either weakly or not significantly correlated with the firing rate (left column of plots). For almost all groups, there was a significant correlation to bursts (middle column of plots). In most of the groups, the proportion of spikes in bursts and those found in sequences were inversely correlated (that is, more sequences were found in cells that showed less bursting). The exception to this were findings in the STN after treatment with MPTP, where high numbers of bursts correlated with higher numbers of sequences. However, this correlation was not significant.

**Figure 4:**
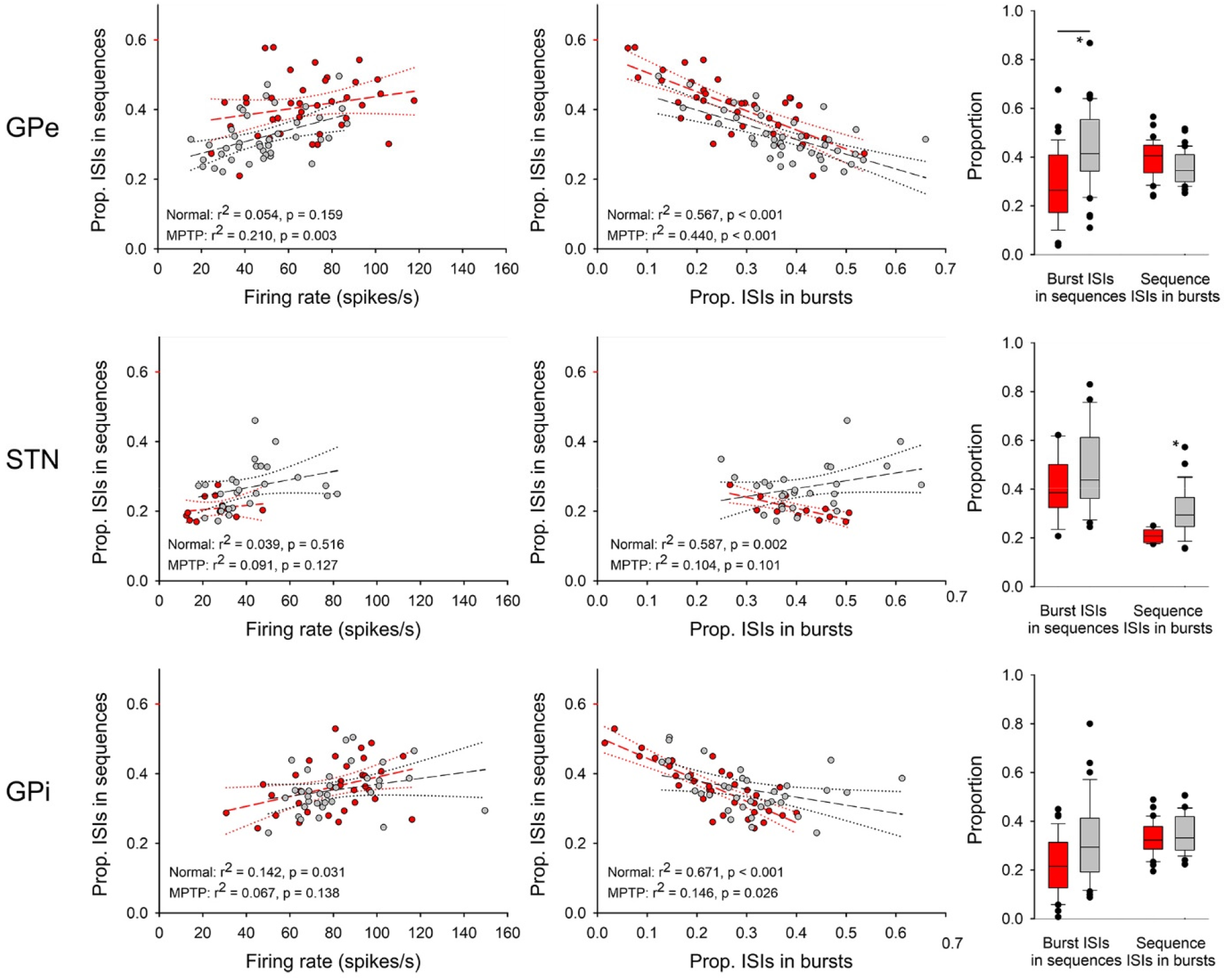
Relation between sequence and bursts. The scatter plots in the left column show the relationship between the proportion of spikes in sequences (ordinate) and the cell’s firing rate (abscissa). The middle column of plots shows the relationship between the proportion of ISIs in sequences (abscissa) and the proportion of ISIs within bursts (ordinate). Each symbol represents a single recorded neuron. The dashed lines represent the result of a linear regression analysis (R^2^ values and p-values are indicated in the figures). The red symbols represent data from the normal state, while the gray symbols represent data from the parkinsonian state. The box plots on the right show the median of the overlap between ISIs that were found to be within sequences and those that were found within bursts. Error bars indicate the 25^th^-75^th^ percentile ranges. The box plots on the left of these figures show the proportion of ISIs within sequences that were also found to be part of a burst. The box plots on the right show the proportion of ISIs within bursts that were also found to be part of sequences. *, p < 0.05; Mann-Whitney test, comparing the data from the normal and parkinsonian states.

The relationship between bursts and sequences was further explored with the analysis shown on the right column of figure 4. The boxplots show the median and inter-quartile range of the proportion of burst ISIs among ISIs that were found in sequences (left box plots) and the proportion of sequence ISIs among those ISIs that were found to be in bursts. Twenty to 40% of ISIs were found in both populations of ISIs. The proportion of burst ISIs among sequence ISIs was increased in the GPe sample in the parkinsonian state, as was the proportion of sequence ISIs among burst ISIs in the STN sample. There were no other significant changes between the normal and parkinsonian states.

## Discussion

The presented results demonstrate that repeating sequences of ISIs are more common than expected by chance in the spontaneous firing of basal ganglia neurons in GPe and STN in the healthy state, and in GPe, GPi and STN in the parkinsonian state. This finding suggests that the formation of such sequences is a feature of neuronal discharge. We also found that the proportion of spikes in sequences changes in GPe and STN in the dopamine-depleted, parkinsonian state, with a reduced probability of finding sequences in GPe and an increased probability of finding them in the STN. The MPTP-treatment did not alter the proportion of spikes in sequences of data collected from GPi. We also show that while there is overlap between the group of ISIs in bursts and those in sequences, these phenomena are not strongly related to one another.

### Parameters that determine sequence detection

Several factors determine the results of sequence detection by the methods used in our study. The length of the recordings is obviously important, with longer recordings favoring the detection of sequences, likely because sequences that occur less frequently can be detected if more time is allowed for their (re-) occurrence. The detection threshold is another important variable in this analysis. We identified a series of ISIs as being a repetition of an earlier ISI series (and thus, a sequence) if all members of the sequence were no more than 1% (0.01) different from the corresponding members of the original sequence. With this criterion, the detection criteria are temporally more stringent for sequences that contain short ISIs, so that results from nuclei with greatly differing average firing rates (such as the STN and the GPi) may not be fully comparable. However, as shown in figure 4 (left column), there was no strong relationship between the average firing rate of neurons and the proportion of spikes in sequences in that neuron (in fact, neurons with a higher firing rate were, if anything, slightly *more* likely to generate sequences than cells with a low firing rate).

The choice of other parameters also substantially influenced the result. As could be expected, the length and the cut-off criterion relation both influenced the likelihood of finding sequences. Sequences in the original data occurred overall more frequently than predicted by chance (see fig. 3, right panels). Further, there is an obvious interaction between the detection of sequences in the original and the shuffled data streams, as ISIs that repeatedly appear in the original data stream would end up being more common in general, thus, favoring random ISI arrangements containing such sequences. Our method of testing whether the original sequence detection was explained by random arrangement of ISIs alone is therefore a conservative estimate.

We also examined how stable estimates of the likelihood of finding spike sequences would be with a given length of ISI stream. Not surprisingly, the use of short data segments resulted in variable estimates, while long data segments provided more stable results (fig. 1B). As a compromise between these results, we used ISI data streams that were 5000 ISIs in length.

### Comparison to literature findings

Precisely replicating patterns of spikes have been described in a variety of other brain locations, mostly belonging to visual or olfactory domains (18, 20-22, 29). As mentioned by other authors, replicating patterns in these systems may present temporal coding of precisely patterned inputs (30), or may represent coded symbols (20, 22). While it cannot be excluded that such temporal or symbolic coding also occurs in the ‘spontaneous’ state, the presence of such patterns in studies of spontaneous discharge (as in our study and others (18) suggests that the patterning of firing is triggered, at least under some circumstances, by (unknown) intrinsic biophysical cellular or network activities.

In sensory systems, repeating ISI patterns in cortical recordings were often improbable when measured against Monte Carlo simulated random arrangements of spikes, as also done in our study (21, 30). We found, however, that the likelihood of finding sequences in spontaneous firing data from the basal ganglia did not reach the same level of significance as those found in the studies of cortical firing. This suggests that external stimuli are represented with less temporal precision in the basal ganglia, or that cellular/network properties of basal ganglia neurons are less conducive to sequence coding than neurons in the cerebral cortex.

In most studies, including ours, the detected sequences do not extend over extended periods. By far, most of the detected patterns consisted of short repetitions (triplets or quadruplets), and, commensurate with the average firing rate of the individual nuclei, the average duration of the detected repeating series of spikes did not extend beyond 25 ms in either segment of the globus pallidus, and 50 ms in the STN.

### Biologic relevance

Between 20 and 40% of all spikes in the GPe, GPi and STN were found to be part of sequences, as defined here. While it seems likely that the tendency to discharge in sequences under rest conditions is related to a given cell’s membrane properties or channel endowment, it is not clear how sequences originate. We did not observe a strong overlap with bursts (sequences could occur within or outside of bursts), thus it is not likely that the two are explained by the same phenomena.

The finding of sequence alterations in the parkinsonian state adds another abnormality to the list of findings that separates parkinsonian from healthy animals. So far, single cell firing in STN, GPe and GPi was found to be characterized by changes in firing rates and the occurrence of (oscillatory and non-oscillatory) bursts (2). The presented findings suggest that, in the parkinsonian state, sequences occur less commonly in GPe, and more commonly in the STN. It is interesting that GPi cells did not show a tendency to produce sequences in our analysis under normal circumstances but did so in the parkinsonian state. The fact that the occurrence of sequences was accentuated in the parkinsonian state may suggest that dopamine loss (or secondary synaptic changes (e.g, ref. (31, 32)) may favor biophysical changes in the networks that incorporate the basal ganglia which may then be conductive to sequence generation.

### Limitations

The most significant limitation of this study is its small sample size. We used the opportunity to analyze data that were available from a different study (see above). Ideally, the analysis would have included a larger number of cells from each structure. Even with these limitations in mind, however, the current study allowed us to show that sequence formation does occur in the extrastriatal basal ganglia, and that the tendency to show sequences is altered by the parkinsonian state.

Another limitation is the fact that we only recorded cells in the spontaneous state. This state was defined by the presence of wakefulness and the absence of overt movements, but this does not rule out that sensory or motor-related information was processed. It would therefore be of interest to study whether sequences become more or less common in animals performing a task, to be able to define the relationship between information processing and sequence formation. It is possible that repeating sequencing are an ‘idling’ phenomenon that would be suppressed when movements are underway. Alternatively, sequences may be part of the information processing currency used by cells to communicate with each other.

Finally, given the prominence of changes in synchrony of firing in the basal ganglia in the parkinsonian state, it would have been interesting to simultaneously record neighboring cells, and to study whether they generate similar or separate sequences. It would be expected that the similarity of sequences in neighboring cells would increase with the onset of parkinsonism. We were not able to test this hypothesis, given the nature of the available data set.

### Summary

To our knowledge, the current report is the first documentation of repeating patterns within series of basal ganglia ISIs and the first time the relationship between parkinsonism (or other disease states) and repeating sequences of neuronal spikes has been explored. The generation of sequences may be an important method of information transfer between these nuclei which, when disturbed, as apparent in the parkinsonian animals, may contribute to behavioral abnormalities. Future delineation of the biophysical mechanisms underlying the generation of sequences in basal ganglia discharge may help us to better define their relevance for the function of this firing behavior under normal and disease states.

## Resource availability

All source data are available at https://doi.org/10.5281/zenodo.6581800. The Matlab codes used to identify sequences and to analyze patterns within the ISI data is available at https://zenodo.org/badge/latestdoi/496319009.

## Acknowledgements

We gratefully acknowledge of Dr. Jesus Soares with some of the initial recording studies. Funding for this analysis was provided by NIH grants P50NS098685 and P50NS123103, and infrastructure grant to the Emory National Primate Research Center (P51OD011132). This research was also funded in part by Aligning Science Across Parkinson’s [ASAP-020572] through the Michael J. Fox Foundation for Parkinson’s Research (MJFF). For the purpose of open access, the author has applied a CC BY public copyright license to all Author Accepted Manuscripts arising from this submission.

---

**Supplementary table.**

|  |  |  | Pooled data (monkeys 1 and 2) |  |  |  | Monkey 1 |  |  |  | Monkey 2 |  |  |  |
| --- | --- | --- | --- | --- | --- | --- | --- | --- | --- | --- | --- | --- | --- | --- |
|  |  |  | Median (IQR) | n | p (comp. to shuffled) | p (MPTP vs normal) | Median (IQR) | n | p (comp. to shuffled) | p (MPTP vs normal) | Median (IQR) | n | p (comp. to shuffled) | p (MPTP vs normal) |
| GPe | Prop. spikes in sequ. | Normal | 0.4176 (0.1036) | 38 | 0.000 | 0.000 | 0.4116 (0.1500) | 13 | 0.000 | 0.005 (38) | 0.4192 (0.0896) | 25 | 0.000 | 0.000 |
|  |  | MPTP | 0.3032 (0.9280) | 40 | 0.000 |  | 0.3132 (0.1168) | 15 | 0.001 |  | 0.2980 (0.0872) | 25 | 0.004 |  |
|  | Prop. time in sequ. | Normal | 0.3289 (0.1242) | 38 | 0.001 | 0.000 | 0.3228 (0.1261) | 13 | 0.000 | 0.003 (34) | 0.3451 (0.0980) | 25 | 0.148 | 0.000 |
|  |  | MPTP | 0.2196 (0.0921) | 40 | 0.868 |  | 0.2286 (0.0860) | 15 | 0.804 |  | 0.2196 (0.1268) | 25 | 1.000 |  |
| STN | Prop. spikes in sequ. | Normal | 0.2016 (0.0296) | 13 | 0.001 | 0.001 | 0.1992 (0.0332) | 12 | 0.002 | 0.000 (18) | 0.2064 | 1 | - | 0.643 |
|  |  | MPTP | 0.2604 (0.0840) | 27 | 0.000 |  | 0.2732 (0.0852) | 14 | 0.001 |  | 0.251 (0.0768) | 13 | 0.002 |  |
|  | Prop. time in sequ. | Normal | 0.1595 (0.3518) | 13 | 0.006 | 0.151 | 0.1595 (0.0579) | 12 | 0.012 | 0.322 (64) | 0.1570 | 1 | - | 0.571 |
|  |  | MPTP | 0.1885 (0.0696) | 27 | 0.002 |  | 0.1885 (0.0551) | 14 | 0.025 |  | 0.1852 (0.0668) | 13 | 0.027 |  |
| GPi | Prop. spikes in sequ. | Normal | 0.3608 (0.1132) | 33 | 0.024 | 0.594 | 0.2932 (0.0736) | 17 | 0.194 | 0.527 (118) | 0.3960 (0.0816) | 16 | 0.064 | 0.093 |
|  |  | MPTP | 0.3460 (0.0720) | 34 | 0.000 |  | 0.3204 (0.0756) | 16 | 0.001 |  | 0.3600 (0.1028) | 18 | 0.000 |  |
|  | Prop. time in sequ. | Normal | 0.3083 (0.1176) | 33 | 0.548 | 0.291 | 0.2497 (0.0671) | 17 | 0.145 | 0.631 (122) | 0.3565 (0.0851) | 16 | 0.632 | 0.064 |
|  |  | MPTP | 0.2772 (0.0721) | 34 | 0.000 |  | 0.2514 (0.0670) | 16 | 0.011 |  | 0.3116 (0.0667) | 18 | 0.004 |  |

Shown are median and inter-quartile range (IQR) of the proportion of spikes in sequences (prop. spikes in sequ.) and the proportion of time in sequences (prop. time in sequ.) in the normal and parkinsonian states, in pooled data from both animals, and separated by animal. Comparisons to shuffled ISI representations were done using the paired Wilcoxon test, while comparison between data from the normal and parkinsonian states were carried out with Mann-Whitney tests.

